# Social agency buffers pain sensitivity during a critical window following spared nerve injury

**DOI:** 10.64898/2026.07.15.738734

**Authors:** Carlee A. Toddes, Kevin N. Bai, Isabel E. Halperin, William R. Keeler, Lukas K. MacMillen, Mitra Heshmati, Sam A. Golden

## Abstract

**Background:** Social buffering of pain, where social contact blunts pain intensity, is well established. However, the degree to which agency over social contact, rather than passive social exposure alone, modifies pain intensity or social behavior is unclear.

**Methods:** Using a social operant self-administration procedure in which mice lever-press for access to their pair-housed partner, we show that voluntary social engagement gates pain after spared nerve injury (SNI). After 8 days of training, mice underwent SNI and then regained access to social self-administration either 1 day (Early) or 5 days (Delayed) following surgery. We compared these mice to groups without social self-administration access, non-contingent social-access, and food self-administration. We assessed changes in pain, or mechanical allodynia, via the von Frey test.

**Results:** Early access to social self-administration after SNI preserved pre-injury social reinforcement and buffered pain. Delayed access similarly preserved social reinforcement but failed to buffer allodynia, revealing an early post-injury critical window for social buffering of pain. Females were more sensitive to both SNI and disrupted social access, while males maintained social reward-seeking after SNI when given early access and showed robust social buffering of pain. Non-contingent (involuntary) social self-administration did not modify pain, nor did food self-administration.

**Conclusions:** There is an early critical window during which voluntary social interactions buffer pain after nerve injury. These results highlight social reward agency and proximity to injury as key variables, since pain outcomes do not generalize to food reward, non-contingent social reward, or delayed access to social reward.

## Introduction

Individual perceptions of pain are highly variable and dependent upon personal history^1^, environment^2^, and behavioral engagement^3^, including day-to-day voluntary social engagement^4^. Social connection provides analgesia^4–6^, and pain frequently precipitates social isolation which in turn enhances pain perception^7–9^. These reciprocal social-pain dynamics substantially alter acute pain trajectories and shape recovery from injury^7,10,11^. Following nerve injury, the motivation to seek and enjoy social interactions can both be disrupted. Maintenance of positive social interactions after injury and social network size predicts improved pain recovery^12^. Notably, atypical pain sensitivity and impaired reciprocal social engagement frequently co-occur in neuropsychiatric conditions with social impairments such as autism spectrum disorder^13–15^. There are few preclinical models to evaluate the impact of social interaction, or social buffering, on pain progression or recovery. Currently available preclinical models do not incorporate social reward-seeking or motivation, a potentially important variable. It remains unclear at which point in the trajectory after an acute painful injury that social motivation is essential for shaping recovery.

Social buffering of pain is reported in both humans^5,6^ and animals^16^. Numerous preclinical reports show overlapping neural circuitry regulate both physical pain and the emotional “pain” of social separation^13,17–22^, suggesting social buffering of pain may occur through these overlapping circuits. Preclinical mouse models evaluating the influence of social behavior on pain reveal reward-related networks as essential for pain processing, pain severity progression^23,24^ and social transfer of analgesia^16^. These studies typically rely on assays such as the three-chamber social test or dyadic social interaction test^4,25^ to evaluate social interactions. These assays use involuntary, passive, or time-constrained social interactions where the experimental animal has few alternatives to interaction. Social tests are often administered at isolated timepoints and historically exclude female mice^26^, offering an incomplete picture of social and sex-specific dynamics. It is well-established that motivation and emotional processing are influenced by voluntary decision-making that is missing in these assays. Social behavioral patterns diverge when mice are given a choice to engage in social interactions via self-administration procedures versus when social exposure occurs via involuntary, non-contingent administration^27,28^. We hypothesize voluntary social self-administration models will help to better define neural mechanisms that underlie social buffering of pain.

Here, we adapt a social self-administration procedure in male and female mice to evaluate the longitudinal influence of voluntary social interactions on pain recovery. We use a well-established spared sciatic nerve injury (SNI) model that produces acute ipsilateral mechanical allodynia that over weeks will progress to chronic neuropathic pain. In this SNI model, pain is worsened by social isolation^29^, and significant social impairment is observed in a modified three-chamber test ∼1 week post-injury^23^ that returns to baseline within a month^25,30^. However, prior studies are temporally constrained and do not directly evaluate whether voluntary social interactions are disrupted by SNI, or whether social interactions may confer ongoing pain buffering.

To address these gaps, mice underwent social self-administration (SSA) in an operant chamber where they lever-pressed for access to freely moving physical interaction with their familiar cage mate^27,31–33^. Next, mice underwent SNI (or sham) surgery to induce neuropathic pain. They were placed back in the operant box for SSA either early on the first post-operative day (POD) or after a delay period of 5 days. This task allowed us to evaluate how the motivational and rewarding features of voluntary social engagement with a familiar partner are modified by neuropathic pain, and reciprocally, how progression of allodynia is shaped by repeated voluntary social interactions. We further examine the temporal dynamics of this relationship by comparing early (POD 1) with delayed (POD 5) return to SSA. We test the necessity of social agency through a non-contingent comparison and evaluate reward-modality specificity through a parallel food self-administration cohort. We demonstrate that voluntary social self-administration in mice confers analgesia within an early critical window after nerve injury.

## Methods and Materials

Procedures common to multiple experiments are summarized below. A detailed description of experimental subjects, apparatus, all behavioral procedures, and the statistical analysis is provided in the Supplemental Methods and Materials. Complete statistical results for every comparison in every figure are reported in Table S2, and sample sizes are reported in Table S1. We describe the specific experiments below. All animal procedures were reviewed and approved by the Institutional Animal Care and Use Committee (IACUC) at the University of Washington. All experiments were performed in accordance with the National Institutes of Health (NIH) *Guide for the Care and Use of Laboratory Animals* and complied with institutional guidelines for the humane care and use of laboratory animals.

### Experiment 1: Temporal profile of voluntary social self-administration after spared nerve injury

The goal of experiment 1 was to determine whether voluntary social self-administration during the acute perioperative window, rather than after allodynia is established, alters the development of mechanical hypersensitivity (Figure 1A). We trained 132 adult male and female C57BL/6J mice on operant social self-administration (SSA) across eight daily discrete-trial sessions. Mice that failed to meet the acquisition criterion (greater than 3 average presses across the last three days of training) were classified as non-acquirers (n=25) and excluded from the primary analyses; their acquisition and lever discrimination data are reported in Figure S1A-B. Mice that directed attack behavior toward the partner during training were removed from the cohort (n=4) as were mice that developed a complete loss of sensation in the back paw following SNI (unresponsive to both von Frey and toe pinch; n=3).

**Figure 1.**
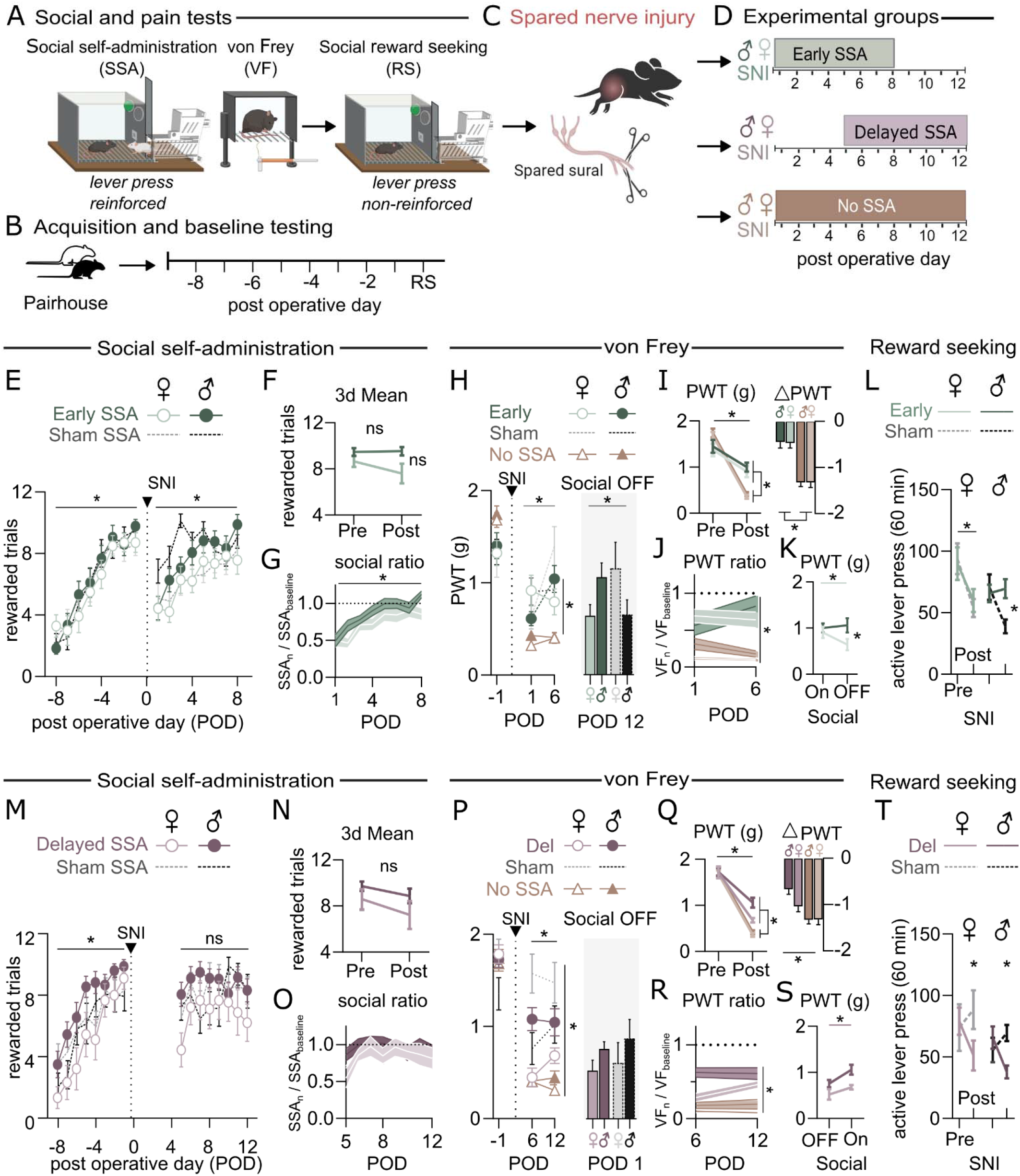
Early but not delayed social self-administration attenuates allodynia following spared nerve injury. (A) Schematic of behavioral procedures: social self-administration (SSA), von Frey testing (VF), and social reward seeking (RS). (B) Timeline of acquisition and testing relative to surgery (POD). (C) SNI surgical preparation. (D) Group design: Early SSA (POD 1; Sham ♀ n=13 ♂ n=15, SNI ♀ n=14 ♂ n=17), Delayed SSA (POD 5; Sham ♀ n=8 ♂ n=9, SNI ♀ n=9 ♂ n=16), No SSA (Sham ♀ n=15 ♂ n=14, SNI ♀ n=16 ♂ n=11). Early cohort (E–L): (E) Rewarded trials across training and post-SNI testing. (F) Three-day mean rewarded trials pre-vs. post-SNI, SNI animals only; no significant group difference in either sex. (G) Social ratio (SSA normalized to baseline). (H) Paw withdrawal threshold (PWT); No-SSA-SNI mice diverge from all other groups at every post-SNI timepoint (*); Early SSA-SNI and Sham remain indistinguishable throughout, including at POD2 after the SSA task ended (right; sig Group × Sex interaction, *, reflecting opposing non-significan1t trends in males and females rather than a group difference in either sex alone). (I) Pre/post PWT and difference scores; No-SSA-SNI shows the largest drop (*). (J) Normalized PWT (von Frey) ratio; No-SSA-SNI significantly lower than all other groups throughout (*). (K) PWT with social access on vs. off, SNI animals only: withdrawing social access significantly increased mechanical sensitivity in females (*) but not males, producing a significant Sex × Condition interaction (*). (L) Active lever presses during reward seeking, pre/post SNI: males show a significant SNI-vs-Sham difference specific to the post-SNI timepoint (*); females show a significant Pre→Post decline irrespective of group (*), with no group difference in either sex. Delayed cohort (M–T): (M–O) SSA acquisition and normalized ratios; comparable to the Early cohort. (P) PWT across PODs; unlike the Early cohort, Delayed SNI mice remain distinguishable from Sham throughout, particularly in females. There is no group prior to the start of social self-administration (right, ns). (Q) Pre/post PWT and difference scores; Delayed SNI intermediate between Sham and No-SSA-SNI (*, both comparisons). (R) Normalized PWT ratio; protection is male-driven (*, males), with a more modest elevation in females (*). (S) PWT with social access off vs. on, SNI animals only: social access significantly decreased mechanical sensitivity in males (*), with a similar but non-significant trend in females, and no significant interaction. (T) Reward seeking pre/post SNI: Delayed SSA-SNI mice showed significantly reduced reward seeking relative to Delayed Sham at the post-SNI timepoint (*). All data mean ± SEM; n’s refer to animals; open symbols female, filled symbols male. Full statistical output (test statistics, df, corrected/uncorrected p values, effect sizes) for every panel is reported in Table S2. *p < 0.05 by Bonferroni-corrected post-hoc test unless otherwise noted.

Mice were assigned to one of three conditions: Early SSA, returned to the operant chambers on post-operative day (POD) 1; Delayed SSA, returned on POD 5, after allodynia onset; or No SSA, received no operant chamber access at any point. Within each condition, mice received either SNI or sham surgery (Early SSA Sham, female n = 13, male n = 15; Early SSA-SNI, female n = 14, male n=17; Delayed SSA Sham, female n = 8, male n = 9; Delayed SSA-SNI, female n = 9, male n = 16; No SSA Sham, female n = 15, male n = 14; No SSA-SNI, female n = 16, male n = 11). All mice, including the No SSA groups, remained pair housed with an age- and sex-matched uninjured partner for the duration of the experiment, so that the groups differed only in access to voluntary social self-administration. This design controls for passive social buffering conferred by co-housing.

Three outcomes were measured: Mechanical sensitivity was assessed by von Frey testing on POD -6, -3, and −1 and averaged within each animal as a Pre-SNI paw withdrawal threshold (PWT), represented in figures as POD -1, and at repeated post-operative timepoints through POD 12. Reinforced social self-administration was assessed daily. Non-reinforced social reward seeking was tested both before SNI (POD –6, -3, and -1, averaged in figures as POD -1) and after the final social self-administration session of each cohort (POD 9 for Early SSA, and POD 13 for Delayed SSA). Mice in the No SSA condition never entered the operant chambers and therefore contributed von Frey data only.

### Experiment 2: Contingent Versus Non-Contingent Social Administration

The goal of experiment 2 was to determine whether the pain buffering observed in experiment 1 requires voluntary control over social access, or whether matched social contact alone is sufficient (Figure 2A). We trained a second cohort of mice (total n = 32) on contingent SSA, using the procedure described in Supplemental Methods, or non-contingent social self-administration procedure in which partner deliveries were matched in number and inter-delivery interval to the mean acquisition performance of the Early SSA cohort in experiment 1, and were delivered independently of lever pressing. Active and inactive lever presses were recorded in the non-contingent group but had no programmed consequence. This design equated the quantity and temporal distribution of social contact across groups and isolated operant agency as the manipulated variable.

**Figure 2.**
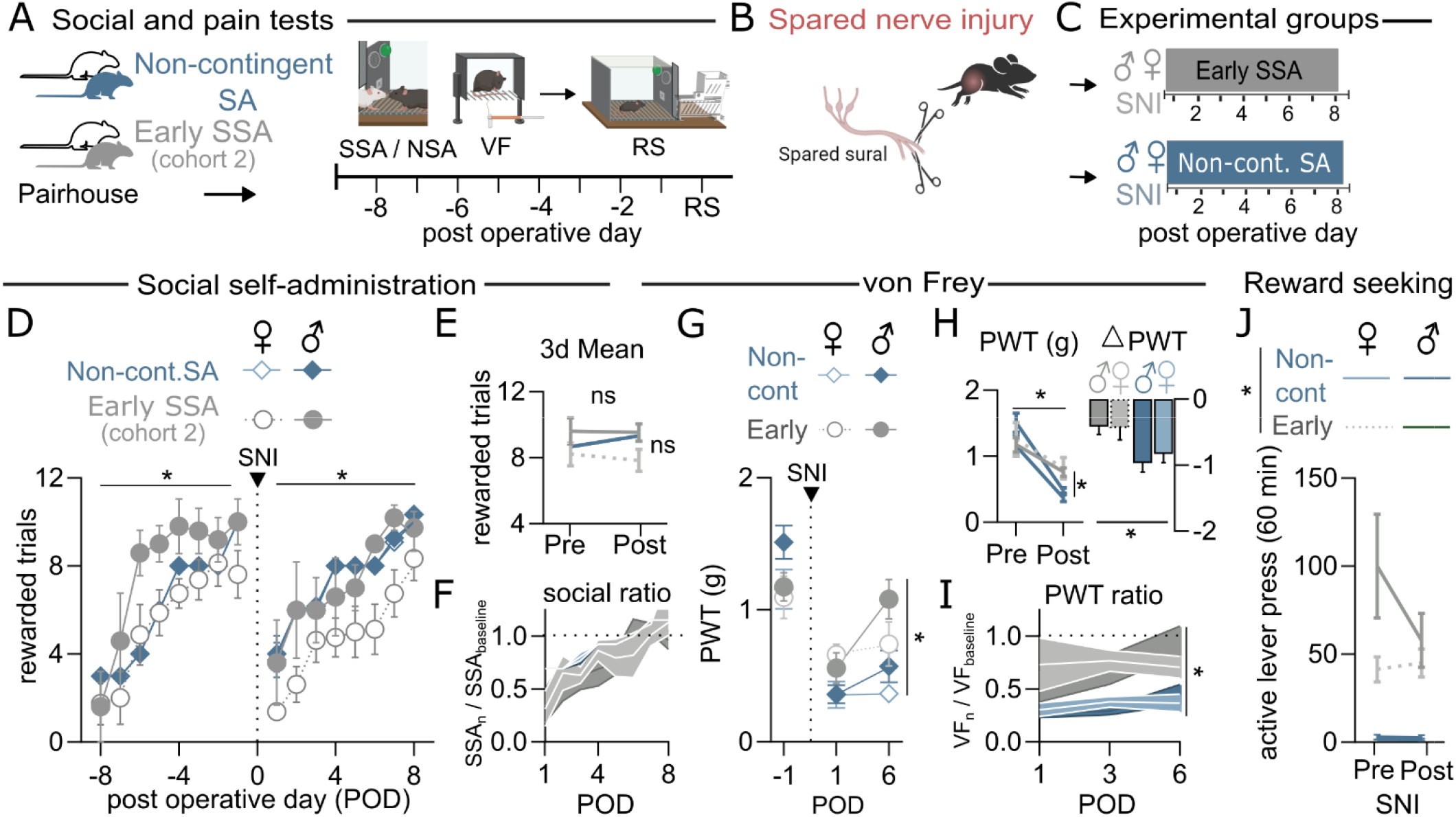
Contingent but not non-contingent social administration attenuates allodynia following spared nerve injury. (A–C) Design schematic and group assignment: Contingent (SNI ♀ n=6 ♂ n=5) vs. Non-contingent SA (SNI ♀ n=11 ♂ n=10), matched for quantity/timing of social contact. (D) Rewarded trials across training and post-SNI sessions. (E) Three-day mean rewarded trials, pre-vs. post-SNI: no significant overall Pre→Post change. (F) Social ratio across POD 1–8; significant effect of Day. (G) PWT across POD –1 to 6; Contingent mice show significantly higher PWT than Non-contingent by POD 6 (*). (H) Pre/post PWT and difference scores; Contingent mice show significantly higher post-SNI PWT and smaller difference scores than Non-contingent (*), no sex effect. (I) Normalized PWT ratio; Contingent significantly higher than Non-contingent overall, most pronounced at Day 6 (*). (J) Reward seeking pre/post SNI; Contingent mice press significantly more than Non-contingent at both timepoints, in both sexes (*). All data mean ± SEM. Diamonds, Non-contingent SA; circles, Contingent SSA. Open symbols female, filled symbols male. Full statistical output in Table S2. *p < 0.05 by Bonferroni-corrected post-hoc test unless otherwise noted.

All mice in experiment 2 received SNI; no sham condition was included, because the relevant comparison is between two SNI groups that differ only in contingency. Mice were returned to their assigned operant context on POD 1 (Contingent SNI, female n = 6, male n = 5; Non-contingent SNI, female n = 11, male n = 10). Von Frey testing was conducted on POD -6, -3, and −1 (averaged within animal as a Pre SNI PWT) and at repeated timepoints through POD 6. Social reward seeking was tested pre- and post-SNI as in experiment 1.

### Experiment 3: Food Self-Administration as a Reward-Modality Control

The goal of experiment 3 was to determine whether the effect of operant engagement on allodynia is specific to social reinforcement or generalizes to a non-social appetitive reinforcer (Figure 3A). We trained a separate cohort of mice (total n = 28) to lever press for highly palatable food pellets under a fixed-ratio 1, 20-s timeout schedule in daily 1-h sessions, with a minimum of eight training sessions before surgery. Mice were not food restricted at any point, so that operant responding reflects the incentive value of the reinforcer rather than caloric need.

**Figure 3.**
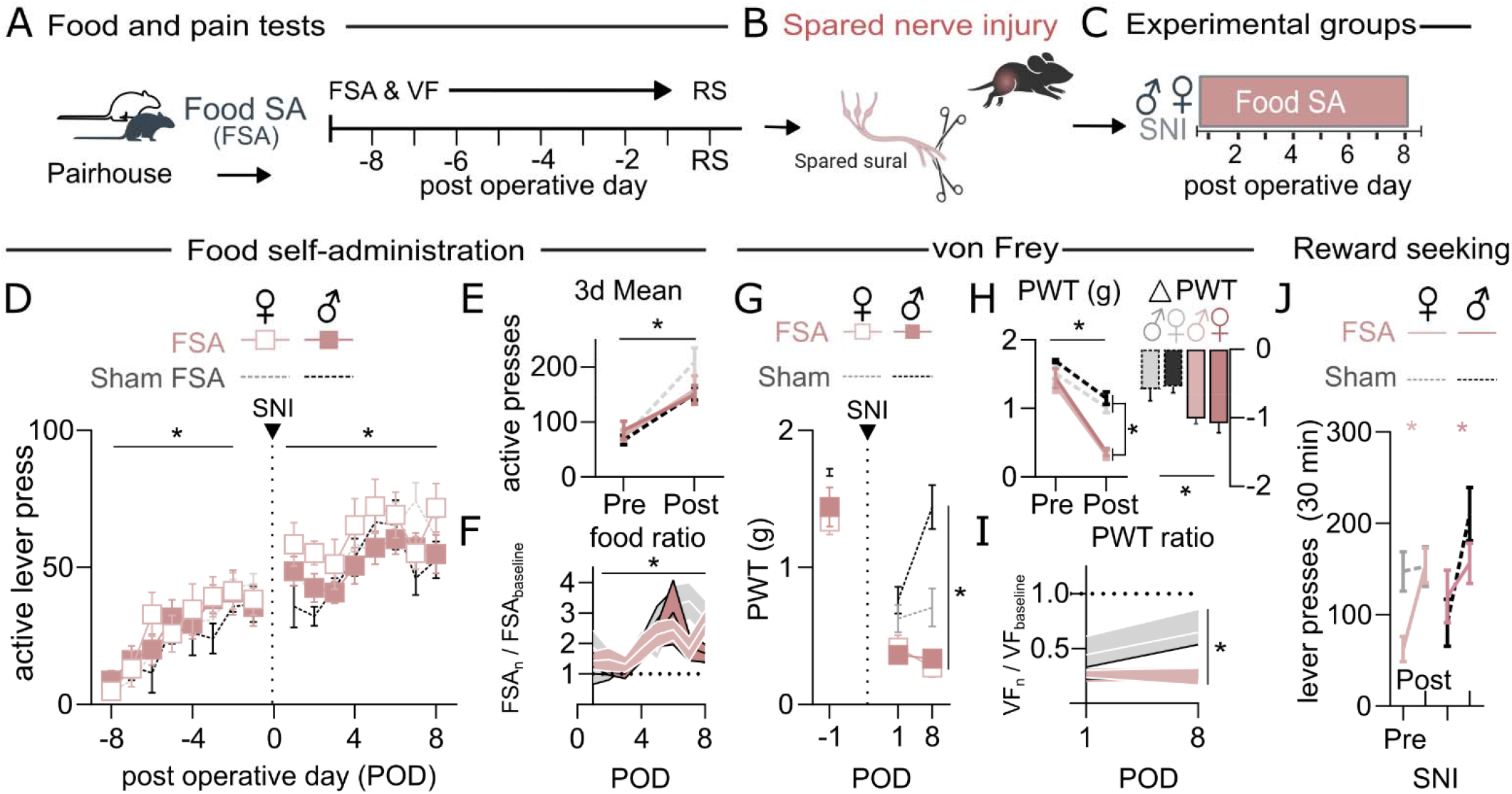
Spared nerve injury increases food self-administration but does not selectively affect food rewrd seeking. (A–C) Design schematic; Food Sham (♀ n=6 ♂ n=7) and Food SNI (♀ n=7 ♂ n=8) returned to FSA chambers at POD 1. (D) Active lever presses across training and post-SNI sessions; significant Group × Day effect in females (*), not in males. (E) Three-day mean; significant Pre→Post increase overall, no Group × Time interaction. (F) Food ratio across POD 1–8; Day effect significant, no group differences. (G) PWT across POD –1 to 8; Food SNI mice (in pink) show significantly lower PWT than Sham mice (in black) (*). (H) Pre/post PWT and difference scores; Food SNI significantly lower post-SNI PWT than Sham (*); no sex difference in difference scores. (I) Normalized PWT ratio; group effect driven by males (*); no group difference in females. (J) Reward seeking pre/post SNI; SNI increases overall food reward-seeking. All data shown as mean ± SEM. Open symbols female, filled symbols male. See Table S2 for detailed statistics. *p < 0.05 by Bonferroni-corrected post-hoc test unless otherwise noted.

Mice then received SNI or sham surgery and were returned to the food self-administration chambers on POD 1 (Food Sham, female n = 6, male n = 7; Food SNI, female n = 7, male n = 8). Von Frey testing was conducted on POD −1 and at repeated timepoints through POD 8, concurrent with food self-administration. Non-reinforced food reward seeking was assessed in a 30-min extinction session pre- and post-SNI.

## Results

### Experiment 1: Social self-administration during an early critical window after SNI buffers allodynia

Mice were trained to acquire social self-administration (SSA) over eight days followed by a social reward seeking task to directly measure motivational salience of social interaction (Figure 1A-B, Video 1). Mice were then separated into either sham or SNI surgical groups (Figure 1C) and returned to the operant chamber at POD 1 or POD 5 to evaluate whether SSA modifies pain behavior at the early or delayed timepoints (Figure 1D). Social housing with familiar conspecifics can buffer pain^34^. To control for passive social buffering, we included a No Social Self-Administration (No SSA) group that were pair housed with same-sex partners but did not undergo the operant task (Figure 1D, Table S1; see Table S2 for detailed statistics). Importantly, all mice (Early SSA, Delayed SSA, No SSA) were pair housed throughout the duration of experiments, with only access to SSA varying between groups.

Mice that acquired SSA and were returned to the social operant chambers on POD 1 (Early group) showed comparable pre-surgical acquisition. Mice returned to their pre-operative baseline of rewarded trials regardless of surgery in both sexes (Figure 1E, Supplemental Figure S1), though females showed lower overall SSA than males (Table S2). Comparison of asymptotic responding (the mean of the last 3 SSA days) before and after surgery revealed no group differences (Figure 1F), and a social ratio (SSA performance normalized to pre-surgical baseline) showed significant increases across SNI days in both sham and SNI groups (Figure 1G). Together, these findings indicate that SNI does not suppress social motivation, as voluntary social access is sustained throughout the post-operative period.

Across repeated von Frey testing (Figure 1H), all groups showed equivalent paw withdrawal thresholds (PWT) prior to surgery. SNI mice that did not receive SSA (No-SSA) exhibited significant pain sensitivity, or reduced PWT, at all von Frey time points. Early SSA-SNI mice were indistinguishable from Shams across all time points. This pattern held when comparing pre-to-post-SNI PWT (Figure 1I) and von Frey ratios (PWT normalized to pre-surgical baseline; Figure 1J). No SSA-SNI mice showed the largest drop in PWT of any group, while the other three groups did not differ. Restricting the analysis to SNI mice, Early SSA-SNI mice showed significantly higher von Frey ratios than No SSA-SNI mice. Male Early SSA-SNI mice also showed persistent allodynia protection following the cessation of social self-administration (Figure 1H, right; Figure 1K), whereas female Early SSA-SNI mice had a decrease in PWT following social self-administration cessation (Figure 1K). Under extinction conditions, *male* Early SSA-SNI mice show significantly higher reward seeking post-SNI compared to Sham controls despite equivalent pre-surgical levels. On the other hand, *female* Early SSA-SNI mice display decreased reward-seeking similar to Sham controls (Figure 1L).

A separate cohort of mice were returned to SSA beginning at POD 5, after allodynia onset. SSA did not differ by group or sex and was equivalent to baseline prior to surgery (Figure M-T, Supplemental Figure S1), confirming that delayed social access did not itself suppress social motivation. However, Delayed SSA-SNI mice showed allodynia and suppressed PWT unlike sham controls. Compared to No SSA-SNI mice, Delayed SSA-SNI mice showed attenuated allodynia (Figure 1P-S), producing a graded pattern (Sham > Delayed SNI>No SSA-SNI) not seen in the Early cohort. This partial pain buffering was primarily male driven. Delayed SNI males showed von Frey ratios near No SSA-SNI levels before social access began (Figure 1P, right), which rose after Day 6 and were sustained through Day 12. Delayed SNI females showed only a modest overall elevation relative to No SSA-SNI females and no significant changes between social self-administration OFF and ON (Figure 1P, 1S).

Unlike the Early social access group, in which male SNI mice showed significantly elevated post-operative reward seeking relative to Sham controls, Delayed group males showed significantly reduced social reward seeking after SNI compared to Delayed SSA Sham mice (Figure 1T, Supplemental Figure S1I). This reversal indicates that the direction of the pain-motivation relationship depends on timing of voluntary social access after nerve injury.

### Experiment 2: Social agency is required for pain buffering

To determine whether social buffering of pain depends on an animal’s agency over seeking social contact, a second cohort was trained on either active SSA (Contingent) or a non-contingent SSA procedure (Non-contingent). Non-contingent social rewards were delivered matched to the average acquisition performance of Experiment 1 mice, independent of lever-pressing (Figure 2A). All experimental mice then received SNI (Figure 2B) and were returned to the operant contexts on POD 1 (Figure 2C). Pre-surgical acquisition and matched post-surgical social contact confirmed that the two groups received equivalent social exposure, different only in contingency (Figure 2D & E).

Both Contingent and Non-contingent groups developed significant allodynia after SNI, but by POD 6 Contingent mice showed significantly higher PWT than Non-contingent mice (Figure 2G). This difference was confirmed by smaller pre-to-post PWT drops and higher von Frey ratios in the Contingent group across sexes (Figure 2H-I). As expected from the operant design, Contingent mice showed dramatically higher social reward seeking than Non-contingent mice both before and after SNI, reflecting active lever pressing in the former and passive receipt in the latter (Figure 2J). This difference was present at both pre- and post-SNI timepoints and did not interact with sex, confirming that the contingency manipulation successfully separated motivation from passive social contact. Together, these results demonstrate that agency in seeking a social partner is necessary for social buffering of pain after SNI.

### Experiment 3: Food self-administration and reward seeking increase following SNI, but do not modify pain

To determine whether mechanical allodynia after SNI is uniquely modulated by social reward, or if self-administration for non-social reinforcers during the early critical window offers a similar buffering effect, we trained a separate cohort to lever press for highly palatable food pellets under non-food restricted conditions (Figure 3A). This cohort received either SNI or sham surgeries (Figure 3B) and were then returned to the operant chambers on POD 1 (Figure 3C).

Food SNI and Sham mice had equivalent active lever presses before and after surgery (Figure 3D). All food-trained mice, regardless of group or sex, showed a robust increase in active lever presses from pre- to post-SNI (Figure 3E-F), with no group differences in the food ratio. Despite comparable self-administration trials as social-trained mice, food self-administration conferred no pain buffering effects. SNI produced a robust and progressive allodynia relative to sham controls in both sexes, evident in raw PWT, pre-to-post difference scores, and von Frey ratios (Figure 3G-I). Food reward seeking under extinction conditions increased significantly from pre- to post-SNI in all mice, though Food SNI mice showed an overall lower increase than Shams that is driven by males (Figure 3J). Taken together, these results indicate that SNI differentially modulates reward seeking depending on the reinforcer. Food self-administration behavior is broadly escalated by SNI, while the effects on social reward seeking depend on timing and contingency of social access. This confirms the analgesic effect of social self-administration is specific to social reinforcement rather than a general consequence of operant reward engagement.

## Discussion

We identify that voluntary operant social self-administration buffers pain as measured by mechanical allodynia in a mouse model of SNI. This buffering effect is (i) temporally bounded, with analgesia depreciating when voluntary social access is delayed post-injury, (ii) contingent on agency, since equivalent non-contingent social contact does not substitute for voluntary engagement in social self-administration, and (iii) specific to social reward, with food self-administration having no effect on allodynia caused by SNI across the same time frame. These findings parallel clinical observations that social connectedness predicts postoperative pain trajectories^35–37^, that decreases in social contacts after injury increase chronification risk^38–40^, and that social presence alone does not reliably affect pain perception^6^. Together these findings indicate the early nerve injury window is a critical period during which voluntary social engagement most effectively buffers pain.

The contrast between the Early and Delayed SSA cohorts defines a critical temporal window. However, our results indicate that even delayed voluntary social interaction has salubrious properties, especially for males. Our results offer a reinterpretation of conflicting reports in the SNI-social literature, where impaired social interaction has been documented at one week but not at later timepoints in socially constrained procedures^25^. Those findings may not reflect spontaneous recovery of social behavior, but rather the closing of an opportunity for social buffering. Once the window has passed, both pain and social motivation outcomes are worse than they would have been with earlier access. This behavior parallels clinical observations of postoperative pain chronification, in which approximately 10% of surgical patients transition from acute pain to persistent neuropathic features within a window during which early social connection may be the most beneficial as an adjuvant^41^. The mechanisms underlying why this window closes are not addressed by the present data. However, previous studies evaluating alterations in allodynia following neuropathic injury have identified adaptations in mesolimbic circuitry, such as those that occur in the nucleus accumbens (NAc) triggered by ventral tegmental area dopaminergic signaling changes^23,24^.

The contingency experiment in Experiment 2 establishes that operant control over social contact is a key factor in social buffering of pain. Non-contingent social delivery, matched in quantity to voluntary acquisition rates, failed to mitigate allodynia. The von Frey trajectory in the non-contingent group resembles mice without any social operant access rather than that of voluntary Early social self-administration mice, ruling out a “social presence is sufficient” interpretation of the buffering effect.

Chronic pain is associated with anhedonia and altered reward processing in both clinical and preclinical settings^25,42–45^. Anhedonia might produce a broad suppression of operant motivation after SNI, but we did not observe this. Food reward seeking was preserved or enhanced after SNI, indicating that the specific decline in social reward seeking does not reflect generalized anhedonia. Food and social rewards differ along multiple dimensions beyond social content, including caloric value and consummatory versus dyadic structure. Social reward may be uniquely vulnerable among reward modalities in the context of neuropathic pain. The present data cannot resolve which specific feature of social reward drives the observed vulnerability but do establish that the effect is not generalizable.

Social buffering of pain via passive social contact was demonstrated in humans and rodents, with the degree of buffering dependent on familiarity^34^. Our data extend these findings by showing that daily, voluntary interactions with an uninjured familiar partner is itself sufficient to mitigate mechanical pain, establishing voluntary social engagement as an effective intervention rather than a byproduct of pharmacological action. Identifying the molecular mechanisms engaged by this behavioral intervention could offer tractable, naturalistic targets for future pharmacological work, positioning social buffering (derived from social agency) as a candidate adjunct to pharmacological standards of care to address dimensions of recovery that traditional analgesics may not directly engage. Where prior preclinical work characterizes social behavior as an output that is altered by pain^3,23,34,46,47^, our data establishes that voluntary social behavior is also an input that shapes the trajectory of pain itself, reinforcing the reciprocal nature of pain and social behavioral domains.

Preclinical literature is historically male-dominated. Testing both sexes throughout this study revealed subtle sex differences in the specific response to social buffering rather than baseline allodynia or motivation. Females appear less motivated to seek social rewards after SNI and showed significantly decreased social buffering in the delayed condition following neuropathic injury. Whether this reflects differences in social reward valuation, descending pain modulation, or recovery kinetics awaits targeted follow-up. Overall, the present data identify three actionable features of social buffering for neuropathic pain: timing, agency, and reward modality specificity. Voluntary social self-administration is a behavioral entry point for continued investigation of the intersectionality of pain and social behavior.

More broadly, the reciprocal relationship we identify between voluntary social engagement and pain responding may extend beyond post-surgical neuropathic pain to neuropsychiatric conditions in which altered pain processing and social impairment frequently co-occur. Autism spectrum disorder is a notable example: atypical nociceptive sensitivity and reduced spontaneous social engagement are both well documented but are rarely studied as interacting phenotypes^13–15^. If agency-dependent social buffering of pain reflects a conserved circuit mechanism, disruption of that mechanism could plausibly contribute to, or result from, altered pain phenotypes in populations with core deficits in social motivation. The operant social self-administration procedure, which dissociates voluntary agency from social contact itself, may be applied in future work examining social and sensory dysfunction across neuropsychiatric disorders, extending this model beyond recovery from nerve injury.

## Acknowledgements

The research was supported by NIDA P30DA048736 (CAT, MH, SAG), NIGMS R35GM146751 (MH), NIDA R01DA059374 (SAG), Washington Research Foundation Postdoctoral Fellowship (CAT), Scan | Design Research Fellowship (CAT), Mistletoe Research Fellowship (CAT), and Weil Neurohub Postdoctoral Fellowship (CAT). Some figures created with BioRender.com.

A version of this manuscript was previously posted as a preprint on bioRxiv (doi: https://doi.org/10.64898/2026.07.15.738734).

Declaration of generative AI and AI-assisted technologies in the writing process: The authors used Claude Sonnet 5, Anthropic for proofreading during the preparation of this manuscript, and for formatting the Supplementary Tables. After using this tool, the authors reviewed and extensively edited the content as needed and take full responsibility for the content of the publication.

## Author Contributions

CAT and SAG conceptualized the study. CAT, MH and SAG wrote the manuscript. CAT, MH and SAG funded the study. CAT, KNB, IEH, WRK, and LKM performed experiments. All authors edited and approved the manuscript.

---

**Supplemental Figure 1.**
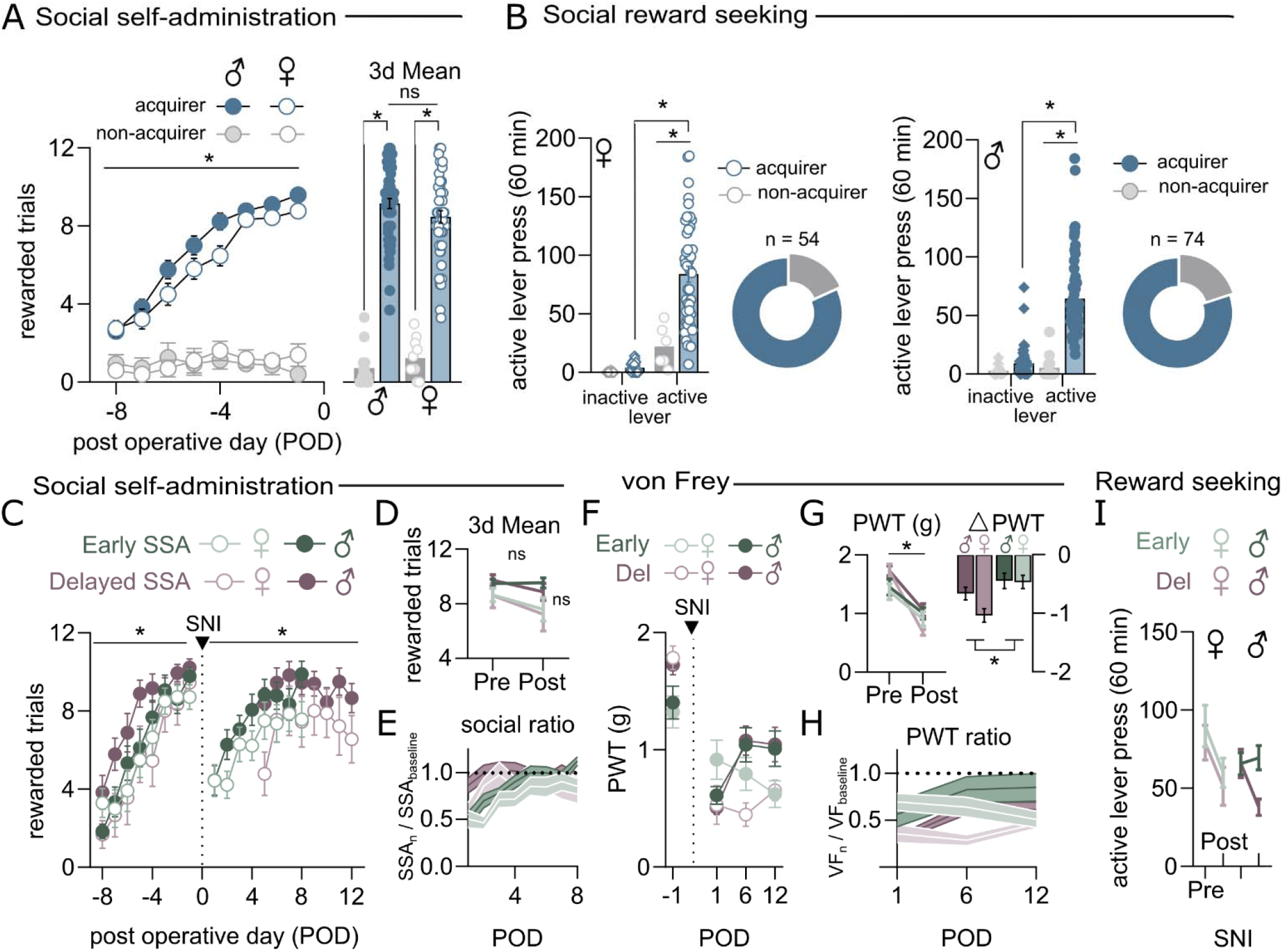
Social self-administration acquisition and reward-seeking behavior across the full Experiment 1 cohort, with acute vs. delayed return to SSA compared after SNI. (A–B) Full cohort (acquirers and non-acquirers), validating lever discrimination prior to surgery. (C–I) SNI-operated acquirers only, Early vs. Delayed SSA, sex as a biological variable throughout. See Table S2 for complete statistical output; group and sex effects are described in the Results and Methods.

