## Supplemental Methods and materials for "Social agency buffers pain sensitivity during a critical window following spared nerve injury"

**Subjects and Housing**

Adult male and female C57BL/6J mice (Jackson Laboratory, stock no. 000664) were used in all experiments. Mice were pair housed with an age- and sex-matched conspecific partner under a 12 h:12 h light/dark cycle with food and water available ad libitum. Mice were bred at the UW vivarium facility. Pair housing occurred at two weeks postweaning and was maintained continuously through the end of each experiment, including for mice assigned to the No Social Self-Administration control condition. Social partners used as reinforcers in the social self-administration procedure were age and sex matched. All behavioral procedures were conducted during the dark phase of the light cycle. Both sexes were included in every experiment, and sex was analyzed as a biological variable throughout. All animal procedures were approved by the Institutional Animal Care and Use Committee and were performed in accordance with the National Institutes of Health Guide for the Care and Use of Laboratory Animals.

**Operant Apparatus**

Mice were trained and tested in Med-Associates operant conditioning chambers (Med Associates Inc., ENV-307A) enclosed within ventilated, sound-attenuating cubicles. Each chamber contained one retractable lever designated active (fixed-ratio 1 reinforced) and one non-retractable lever designated inactive, all positioned 2.4 cm above the stainless-steel grid floor. A custom 3D-printed mouse delivery device^1^ was attached to the chamber wall adjacent to the active lever and housed the social partner between trials. The partner was admitted to the chamber through an automatic guillotine-style door. Reinforcement was signaled by a 2-s tone cue (2900 Hz, 20 dB above background). Chamber events were controlled and recorded using MED-PC V software.

**Social Self-Administration**

The operant social self-administration (SSA) procedure was adapted from previously published designs^1,2^ and is shown schematically in Figure 1A.

**Magazine training.** Prior to operant training, mice received three 5-min magazine training sessions separated by 15-min inter-session intervals. During each session the social partner-paired house-light was illuminated, the tone cue was presented, and the social partner was immediately inserted into the chamber. Levers were not available during magazine training.

**Acquisition training.** Acquisition training consisted of eight daily 48-min sessions using a discrete-trial design^1–3^. Each session comprised twelve 4-min trials. Each trial was initiated by house light illumination, followed 10 s later by insertion of the active lever. Mice had 60 s to satisfy the fixed-ratio 1 requirement. A successful press retracted the lever, presented the 2-s conditioned tone cue, and opened the guillotine door for 120 s of social partner access. Each trial included a 120-s inter-trial interval during which the house light was extinguished and levers were retracted. Trials on which the mouse failed to press within the 60-s response window were scored as omissions and terminated without partner delivery. Active lever presses, inactive lever presses, rewarded trials, and latency to press were recorded for every session.

**Acquisition criterion and exclusions.** Mice were classified as acquirers if they pressed an average of three times across the last three days of training. Mice that did not meet this criterion were classified as non-acquirers and were excluded from all primary analyses. Acquisition and lever discrimination data for the full Experiment 1 cohort, including both acquirers and non-acquirers, are presented in Figure S1A-B. Of 130 mice trained in Experiment 1, 103 met the acquisition criterion and 27 did not. Mice that directed attack behavior toward the social partner at any point during training were removed from the cohort (4 mice across all experiments).

**Social reward seeking.** Non-reinforced social reward seeking was assessed in 60-min extinction sessions^1,3^. The house light and active lever were presented as during training, but active lever presses produced only the conditioned tone cue, delivered on a 20-s fixed-interval schedule, and were never followed by social partner delivery. Cue presentations, active lever presses, and inactive lever presses were recorded. Reward seeking was tested on POD −1 in all social cohorts and again the day after the last social self-administration session.

**Non-contingent Social Administration**

Mice assigned to the non-contingent condition received social partner deliveries that were matched in number and inter-delivery interval to the mean acquisition performance of the Early SSA cohort in Experiment 1. Partner deliveries occurred on this predetermined schedule regardless of the animal's behavior. The house light, tone cue, and door opening were presented exactly as in the contingent procedure, and the duration of partner access was identical. Both levers were present and both active and inactive lever presses were recorded, but neither had any programmed consequence for partner delivery. Session length, number of sessions, and handling were identical to the contingent procedure.

This design equates the total quantity, duration, and temporal distribution of social contact between groups, and decouples social contact from voluntary operant control. It therefore isolates agency over social access, rather than social exposure per se, as the manipulated variable.

**No Social Self-Administration Control**

Mice assigned to the No Social Self-Administration (No SSA) condition in Experiment 1 were pair housed with age- and sex-matched uninjured partners throughout the experiment but never received access to the operant chambers, either before or after surgery. These mice underwent SNI or sham surgery and von Frey testing on the same schedule as the SSA groups. Because pair housing was maintained continuously in all conditions, the No SSA groups control for passive social buffering and social enrichment conferred by cohousing^4^, isolating operant social self-administration as the only variable that differed between conditions. No SSA mice were transported and handled on the same schedule as SSA mice to equate handling stress.

**Food Self-Administration and Food Reward Seeking**

The food self-administration procedure followed our previously published protocol^5^. Mice received one 30-min magazine training session in which 15 food pellets (20 mg; TestDiet catalog no. 1811142; 12.7% fat, 66.7% carbohydrate, 20.6% protein) were delivered on a fixed schedule paired with a 2-s cue light. Operant training consisted of daily 1-h sessions under a fixed-ratio 1 schedule with a 20-s timeout. Each session began with house light onset, followed 10 s later by insertion of the central retractable active lever. Active lever presses illuminated the food-paired conditioned stimulus for 2 s and delivered one food pellet; presses during the 20-s timeout were recorded but had no programmed consequence. Inactive lever presses had no programmed consequence. Mice received a minimum of eight training sessions before surgery. Active lever presses, inactive lever presses, and head entries into the food port were recorded.

Mice were not food restricted at any point in the experiment, so that operant responding reflected the incentive value of the palatable reinforcer rather than caloric need. Non-reinforced food reward seeking was assessed in 30-min extinction sessions in which active lever presses produced the food-paired cue alone, without pellet delivery.

**Spared Nerve Injury**

The spared nerve injury (SNI) model was performed as previously described^6,7^. Mice were anesthetized with 1.5 to 2.5% isoflurane in a 30% N₂O / 70% O₂ mixture and maintained on a heating pad. The left sciatic nerve was exposed at the level of its trifurcation into the sural, tibial, and common peroneal branches. The tibial and common peroneal nerves were tightly ligated with 7-0 silk suture and severed distal to the ligation, leaving the sural nerve intact and unmanipulated. Muscle and skin were closed in layers with sutures. Sham mice underwent identical anesthesia and surgical exposure of the sciatic nerve without ligation or transection of any branch. All sham and SNI mice received perioperative Ketoprofen (5mg/kg,sc) which has a working time of 12hrs. Operant testing for the Early cohort began approximately 20-24hr hrs following surgery and ketoprofen perioperative injection. POD D1 SNI for all groups occurred approximately 27hrs after ketoprofen administration. Mice were allowed to recover alone and return to righting (approximately 20 minutes) before return to the home cage.

**Von Frey Mechanical Sensitivity Testing**

Mechanical sensitivity was assessed using calibrated von Frey monofilaments (Stoelting Co.) applied to the plantar surface of the ipsilateral (left) hind paw, within the sural nerve territory. Immediately following self-administration sessions, mice were placed individually in Plexiglas enclosures on an elevated wire mesh platform and allowed to habituate for 45 to 60 min. Filaments were applied using the ascending stimulus method^8,9^, in which filaments were presented in ascending order of force, each applied perpendicular to the plantar surface with enough force to cause slight bending and held for a maximum of 2 s. Each filament was applied 10 times with an inter-application interval of 5-s. Brisk paw withdrawal, flinching, or licking of the stimulated paw was scored as a positive response. PWT was defined as the maximal response threshold in grams, that is, the lowest filament force that reliably elicited a positive response^72^. Filament series spanned 0.016 – 2g. Testing was conducted by experimenters blind to surgical condition. Von Frey testing was conducted on PODs -6, -3, and −1 and on PODs 1, 3, 6, 9, 12 in Experiment 1, PODs -6 through 6 in Experiment 2, and PODs −6 through 8 in Experiment 3.

**Derived Measures**

Several normalized measures are reported alongside raw values.

• **Social ratio** was calculated as rewarded trials on a given post-operative day divided by the mean rewarded trials across the final three pre-surgical baseline sessions (SSA_n / SSA_baseline).
• **Food ratio** was calculated identically, using active lever presses in the food self-administration procedure (FSA_n / FSA_baseline).
• **Von Frey (VF) ratio** was calculated as PWT on a given post-operative day divided by the pre-surgical average PWT measured on POD -6, -3, and −1 (VF_n / VF_baseline).
• **Difference scores** were calculated as post-SNI minus pre-SNI PWT, where the post-SNI value is the mean PWT during social self-administration. Early: POD 1, 3, 6, 9; Delayed: POD 6, 9, 12; Contingent/Non-contingent: POD 1 & 6; Food: POD 1 & 8. The pre-SNI value for all groups and experiments is the average of POD -6, -3, and−1 baselines.
• **Social ON scores** were calculated as the average PWT taken from von Frey tests that occurred concurrent with social self-administration. Early: POD 1, 3, 6, 9; Delayed: POD 6, 9, 12. Whereas **social OFF** was calculated as the average PWT on days following the cessation of social self-administration (Early; POD 12) or prior to the start of social self-administration (Delayed; POD 1 & 3)
• **Three-day means** for self-administration measures were calculated across the last three pre-surgical sessions and the first three post-surgical sessions.

**Statistical Analysis**

Statistical analyses were performed in Python (v3.12) using the pandas, pingouin (v0.6.1), and SciPy libraries. All data are presented as mean ± SEM. Sample sizes for each experiment are reported in the corresponding figure legend and in Table S1. Complete statistical results for every comparison in every figure, including test statistics, degrees of freedom, uncorrected and corrected p values, and effect sizes, are reported in Table S2.

Repeated measures collected across post-operative days or timepoints, including rewarded trials, social and food self-administration ratios, paw withdrawal threshold, and normalized von Frey ratio, were analyzed using mixed-model analyses of variance (ANOVAs) with Group and Sex as between-subjects factors and Day, Time, or Timepoint as a within-subjects factor. Where a significant Group × Sex × Day or Group × Sex × Time interaction was observed, follow-up mixed ANOVAs were performed separately within each sex, with Group as the sole between-subjects factor. Mauchly's test was used to assess sphericity for all within-subjects factors with more than two levels, and Greenhouse-Geisser-corrected p values are reported wherever sphericity was violated.

Difference scores (post-SNI minus pre-SNI PWT) were analyzed using two-way ANOVAs with Group and Sex as between-subjects factors. Acquisition and active-versus-inactive lever discrimination among acquirers and non-acquirers (Figure S1A, B) were analyzed using one-way ANOVAs; equality of variance was assessed using the Brown-Forsythe and Bartlett tests.

Significant omnibus effects were followed by pairwise comparisons corrected for multiple comparisons using the Bonferroni method: paired t tests for within-subject comparisons across days or timepoints, and independent-samples t tests for between-group comparisons. Post hoc comparisons following one-way ANOVAs (Figure S1A, B) used the Tukey multiple-comparisons test. Effect sizes are reported as partial eta-squared (η²p) for ANOVA main effects and interactions, R² for one-way ANOVAs, and Hedges' g for pairwise comparisons.

Statistical significance was set at α = 0.05 and is denoted in figures as *p < 0.05; n.s., not significant. All tests were two-tailed. No formal power analysis was conducted to determine sample size; group sizes were based on prior work.
