## Supplementary material for "Social agency buffers pain sensitivity during a critical window following spared nerve injury": Table S1

### Table S1. Animal cohort details and group sizes

*n = individual animals. Exclusion criteria: failure to acquire SSA criterion, observation of attack behavior, or missing post-surgical VF data. No SSA animals were pair-housed controls without operant chamber access. Food SA cohort maintained separately (see main text).*

| Experiment | Group | Sex | Surgery | n (enrolled) | Exclusions (n) | Exclusion Reason | Final n (analysis) | Measures Included |
|---|---|---|---|---|---|---|---|---|
| **Experiment 1: Acute social self-administration (SSA) + No SSA controls** | | | | | | | | |
| | Acute SSA | Female | Sham | 13 | 0 | — | 13 | SSA, Von Frey, Reward Seeking |
| | Acute SSA | Male | Sham | 15 | 0 | — | 15 | SSA, Von Frey, Reward Seeking |
| | Acute SSA | Female | SNI | 14 | 0 | — | 14 | SSA, Von Frey, Reward Seeking |
| | Acute SSA | Male | SNI | 17 | 1 | Missing post-SNI VF data (M11, SSA_Cohort3) | 16 | SSA, Von Frey, Reward Seeking |
| **Experiment 1 Total** | | | | **59** | **1** | | **114** | |
| **Experiment 1b: Delayed SSA (POD 5) + No SSA controls (shared)** | | | | | | | | |
| | Delayed SSA | Female | Sham | 8 | 0 | — | 8 | SSA, Von Frey, Reward Seeking |
| | Delayed SSA | Male | Sham | 9 | 0 | — | 9 | SSA, Von Frey, Reward Seeking |
| | Delayed SSA | Female | SNI | 9 | 0 | — | 9 | SSA, Von Frey, Reward Seeking |
| | Delayed SSA | Male | SNI | 18 | 2 | Missing post-SNI VF data (M56 SSA8; M05 SSA_Cohort5) | 16 | SSA, Von Frey, Reward Seeking |
| **Experiment 1b Total** | | | | **44** | **2** | | **98** | |
| **Experiment 1: Pairhoused Controls** | | | | | | | | |
| | No SSA (pair-housed control) | Female | Sham | 15 | 0 | — | 15 | Von Frey only |
| | No SSA (pair-housed control) | Male | Sham | 14 | 0 | — | 14 | Von Frey only |
| | No SSA (pair-housed control) | Female | SNI | 16 | 0 | — | 16 | Von Frey only |
| | No SSA (pair-housed control) | Male | SNI | 11 | 0 | — | 11 | Von Frey only |
| **Experiment 1 Controls Total** | | | | **56** | **0** | | | |
| **Experiment 2: Contingent vs. Non-contingent social access (SNI only)** | | | | | | | | |
| | Contingent (Acute SSA cohort 2) | Female | SNI | 6 | 0 | — | 6 | SSA, Von Frey, Reward Seeking |
| | Contingent (Acute SSA cohort 2) | Male | SNI | 5 | 0 | — | 5 | SSA, Von Frey, Reward Seeking |
| | Non-contingent (yoked) | Female | SNI | 11 | 0 | — | 11 | Von Frey, Reward Seeking (lever presses only) |
| | Non-contingent (yoked) | Male | SNI | 10 | 0 | — | 10 | Von Frey, Reward Seeking (lever presses only) |
| **Experiment 2 Total** | | | | **32** | **0** | | **32** | |
| **Experiment 3: Food self-administration (FSA; separate cohort)** | | | | | | | | |
| | Food SA | Female | Sham | 7 | 0 | See note | 7 | FSA, Von Frey, Reward Seeking |
| | Food SA | Male | Sham | 6 | 0 | See note | 6 | FSA, Von Frey, Reward Seeking |
| | Food SA | Female | SNI | 8 | 0 | See note | 8 | FSA, Von Frey, Reward Seeking |
| | Food SA | Male | SNI | 8 | 0 | See note | 8 | FSA, Von Frey, Reward Seeking |
| **Experiment 3 Total** | | | | **29** | **0** | | **29** | |
